# A Biological Signature of Quantum Gravity?

**DOI:** 10.1101/2024.09.25.614787

**Authors:** Irfan Lone

## Abstract

In a recent proposal on the experimental tests of quantum gravity creation of non-Gaussianity in a Bose-Einstein condensate (BEC) has been suggested as a decisive confirmation of quantum gravity. In a related proposal, a gas of ultracold Rb or Cs atoms has previously been suggested as a possible platform for tests of quantum gravity. Since a practical demonstration of above proposals is a very challenging and costly affair, exploring cost-effective alternatives to these technologically demanding experimental protocols becomes very important. We here show that the phenomenon of Bicoid (Bcd) gradient formation in the early fruit fly embryo, considered basically here as a multipartite quantum system with an ensemble of initial states and a unitary evolution *U* that implements a quantum Newtonian Hamiltonian over this gravitationally interacting system, naturally combines the essential features of above proposals in a single system giving a viable signature of quantum gravity through the creation of non-Gaussianity. We conclude that although the phenomenon of Bcd gradient formation in the early Drosophila embryo is accompanied by quantum gravitational effects, it might need further experiments to verify such a noval claim.

---

The existing canonical theories of quantum gravity, like loop quantum gravity [1–3] and string theory [4–7], have thus far remained empirically unsubstantiated. Recent experimental observations of GRBs [8] have only placed more stringent bounds on the observability of such effects as violations of Lorentz symmetry predicted by certain theories of quantum gravity [9]. Therefore, non-relativistic or low-energy quantum gravity models have received renewed interest in recent years [10–13]. This resurgence of interest has also occurred largely because of the prospect of experimental tests made feasible by new developments in technology. Such indirect and phenomenological approaches, in addition to potentially resolving the question of the quantum nature, or otherwise, of gravity, are also needed, for example, to assess the promise of future experiments on quantum gravity [12]. In this regard, an important scale where gravitational effects of massive quantum systems become relevant is the Planck mass scale, whereby, making use of the natural units, (*c* = *ħ* = 1), all physical quantities can be expressed in terms of the Planck mass,

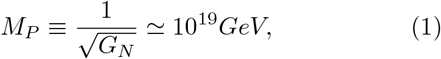

where *G*_*N*_ denotes the Newton’s constant of universal gravitation [14]. This mass scale, it is thought, shall be soon within reach in laboratory settings, thanks to the rapidly developing field of quantum information science.

Therefore, in a recent proposal on the experimental tests of quantum gravity it has been suggested that the creation of non-Gaussianity in the quantum field state of matter of a Bose-Einstein condensate (BEC) can be taken as a decisive confirmation of the existence of quantum gravitational interactions between its constituents [15]. This is one of the most promising proposals, among various suggestions that have been put forward in recent years[16–24], that relies on the gravitational effects of massive quantum systems. A single spherical BEC of∼ 10^9^ atoms and radius∼ 200*µm* is left to evolve and self-interact gravitationally for about *t* ∼ 2*s* and the creation of non-Gaussianity in the system is taken as a signature of quantum gravity. In a related proposal, a test of quantum gravity using a gas of ultracold Rb or Cs atoms has also been suggested previously [25]. The main advantage in using quantum gases, in contrast to the entanglement-based proposals [16, 17], is that through the use of external magnetic or optical fields the noise from the electromagnetic interactions can be suppressed without affecting the strength of the sought gravitational interactions between its constituents [15]. Crucially, in both proposals interactions mediated via a quantum-valued gravitational field provide a signature that is distinctly different from classical gravitational interactions for appropriate initial states of the system.

Nevertheless, a practical demonstration of above proposals would require very sensitive laboratory apparatus to create and preserve the desired quantum states, apart from being potentially very costly. Furthermore, since many of these proposed experiments rely on entangle-(1) ment as a witness of quantum gravity, it has been argued that such effects could also be generated through a classical gravitational field [26]. See [27], however. In view of these limitations, exploring cost-effective and water-tight alternatives to expensive and technologically demanding experimental protocols becomes very important. An area that has in recent years seen significant activity along similar lines is electroweak quantum chemistry [28, 29].

There has been an increasing interest in exploiting the optical properties of certain biological molecules called amino acids to verify the predictions of standard model, offering a potentially noval testing ground for the low energy scale of particle physics [28, 29]. Similarly, recent technological advances in controlling and manipulating fluids have been suggested as possible platforms for analogue simulations of quantum gravity [30]. Here a flowing fluid provides an effective curved spacetime allowing the simulation of gravitational geometries and other related phenomena. More recently, spin nematics have allowed the simulation of gravitational waves presenting a potential opportunity to carry out experiments otherwise thought to be impossible with traditional gravitational wave interferometers like LIGO and the Virgo [31]. Such noval studies not only deepen our understanding of fundamental physics but also showcase the power of integrating otherwise unrelated scientific disciplines.

In this Letter we show that the phenomenon of Bicoid (Bcd) morphogen gradient formation in the early fruit fly embryo naturally combines the essential features of the Bose gas [25] and the BEC proposal [15] in a single system, yielding a biological signature of quantum gravity.

A morphogen gradient is defined as a concentration field of a molecule that acts as a dose-dependent regulator of cell differentiation [32]. The prototypical morphogen gradient is that of the transcription factor Bicoid (Bcd), formed in the early Drosophila melanogaster, fruit fly, embryo and plays a key role in the determination of embryonal axis of the organism [33–39]. Recent experiments, utilising fluorescence correlation spectroscopy (FCS), have revealed multiple modes of Bcd transport at different spatial and temporal locations across the embryo [40]. These observations have been explained through a model fundamentally based on quantum mechanics [33–35]. This has been achieved by making use of an aspect of quantum information theory called a discrete time quantum walk, in which a Bcd molecule can act as a qubit due to chirality superposition states (Fig. 1) [41]. In such a treatment the degradation of Bcd is modelled as a unitary noise that is intrinsic and that does not cause any entanglement with the environment, so that the system remains in an essentially pure state during the course of its evolution [33–35]. It thus acts like a classical fluctuating field such that the dynamics of the system is unitary, yet stochastic [42].

**FIG 1.**
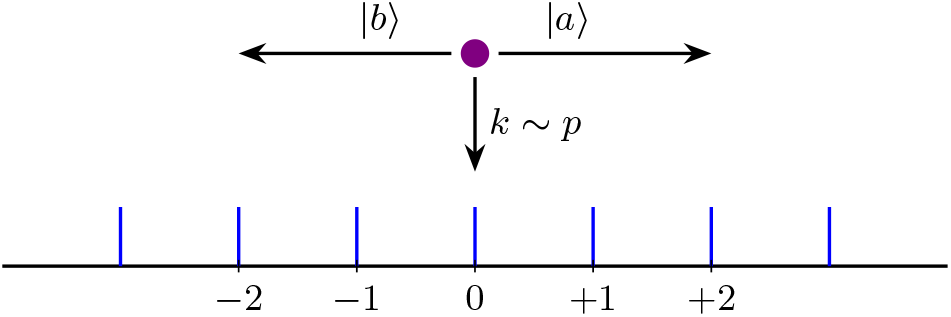
Schematic of the quantum-classical model. A Bcd molecule (shown by the violet colored dot) can, respectively, make a transition to the right or to the left depending on its chirality state of |*a* ⟩ or |*b* ⟩ or it can get degraded at a rate *k*.

Thus, the state of the system at any given time in such a model is represented by a vector Ψ in a finite-dimensional Hilbert space ℋ. Molecules are represented, just like in particle-based reaction–diffusion schemes [43], by point particles undergoing a Markovian quantum diffusion process in presence of a unitary noise that introduces stochasticity into their dynamics. The basic hypothesis underlying this treatment is that transient quantum coherences in collaboration with unitary noise are responsible for the observed dynamics and relaxation to a non-equilibrium steady-state of the Bcd morphogen gradient [44]. It has thus been shown that the multiple dynamic modes of the Bcd gradient are a consequence of a quantum-classical dynamics in which transient quantum coherences play an essential role. Therefore, the quantization of Bcd system accounts for its relaxation to a non-equilibrium steady-state. Such a system may be described, alternatively, by a non-relativistic scalar quantum field 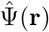 that creates a particle at position **r**, such that *â*^*†*^ and *â* are the creation and annihilation operators of this field [15]. In this way one is also able to account for the particle number fluctuations in the system [45].

The Bcd system actually belongs to a class of closed quantum systems that have to be made ergodic by the addition of a small random perturbation [49–51]. We provide more details about such systems in the Supplementary Information [44]. Addition of random unitary noise to the Bcd system thus relaxes the underlying quantum dynamics to statistical mechanics [33–35]. It will be argued here that the quantum gravitational interaction between the molecules in the system causes the emergence of the observed non-Gaussian distribution. The model that one must therefore consider is the following

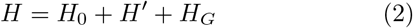

where *H*_0_ represents the free Hamiltonian of Bcd molecules, assumed here to be like a Bose gas of microscopic particles,

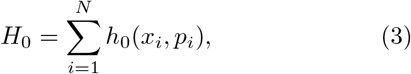

and *H*^*′*^ is the perturbation due to the unitary noise that leads to statistical mechanics in the long time limit. The resulting perturbed Hamiltonian is still Hermitian and therefore generates time evolution operators that are unitary, but now include a stochastic part as well. Formally, one can write such an operator as

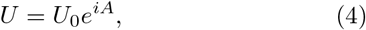

where *A* is a stochastic Hermitian operator determined by a given perturbation and *U*_0_ is the time evolution operator for the original quantum process in absence of perturbation [52]. The Hamiltonian *H*_*G*_ in Eq. (2) accounts for the quantum-valued gravitational interaction in the system. It is this last term that will be our main concern in the present paper.

The Bcd morphogen inside the embryonic syncytium may now be considered as an ensemble of Bose particles with their two chirality states denoted as |*a* ⟩ and |*b* ⟩, and thus viewed as a continuous-variable bosonic quantum system [53]. In other words, a concentration field of Bcd molecules is assumed here, for all practical purposes, to behave as a dilute Bose gas of structureless particles for a very brief instant of time (∼ 2*s*), sufficient for a quantum-valued gravitational interaction between them [15, 25]. In the language of quantum information theory, we have a multipartite quantum system and an ensemble of initial states with a unitary evolution *U* that implements a quantum Newtonian Hamiltonian over this gravitationally interacting system [24]. See Supplementary Information, where we use recent results on LOCC criterion and swapping to argue that such a claim should be testable experimentally. One may then identify the similarity of the Bcd dynamics [33–35] with the Bose gas based proposal [25] in the following manner.

In the Bose gas proposal each atom is essentially prepared in a given initial state denoted |*a* ⟩, before being placed in an equal coherent superposition of states |*a* ⟩ and |*b* ⟩ at *t* = 0. These two modes are then allowed to interact gravitationally for some finite period of time and the system is then subjected to a measurement process [25]. In the Bcd system each molecule is initially produced naturally in a given chirality state, predominantly the left, denoted here also as |*a* ⟩ and |*b* ⟩. The coherent superposition of the two chirality states of the system is then affected through the application of a Hadamard transform applied at *t* = 0 [33–35]. The particles then evolve and may briefly interact gravitationally before being subjected to a degradation process that can be modelled as a unitary noise, which is completely equivalent to a measurement performed on the system [33–35]. It is the gravitational interaction between the molecules that causes the emergence of non-Gaussianity in the system as the experiments based on lattice light sheet microscopy attest [54]. The role of degradation is to introduce stochasticity into the dynamics of the system.

The contribution to the Hamiltonian due to the direct gravitational interaction between the particles will then be given by

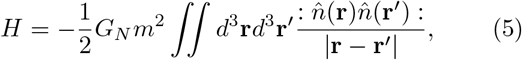

where *m* denotes the mass of each Bcd particle and *G*_*N*_ is Newton’s constant of universal gravitation as already stated in the beginning. The operator

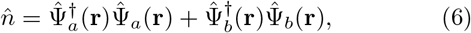

represents the total particle density and :: denotes normal ordering [25]. 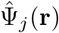 is the standard bosonic field operator that annihilates a particle from state |*j* ⟩ at point **r**, and obeys the standard commutation relations

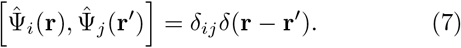

Furthermore, in writing Eq. (5) one is implicitly assuming that the gravitational field can be quantum valued. More specifically, Eq. (5) is re-written as

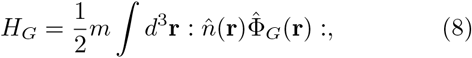

Where

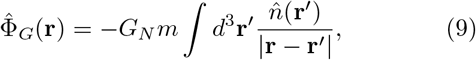

is the quantum valued gravitational potential due to source masses at location **r**^*′*^ [25].

Bcd is initially produced in a predominantly left-hand chirality state |*b* ⟩ through enzymatic intervention, before being placed in a coherent superposition of |*a* ⟩and |*b* ⟩ states. After evolving under the effects of the unitary noise for time *t*, the molecules relax to a non-equilibrium steady-state characterised by a non-Gaussian distribution. For a pure state the density operator is given by

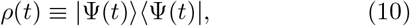

with the normalization

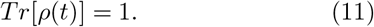

The effect of the unitary noise during the whole process is equivalent to a measurement performed on the system [33–35]. Since the density operator for the system is given by Eq. (10), one may consider the decoherence in the form of randomly occurring projective measurements performed on the system in order to simulate its probability distribution. This, it can be shown [55], is completely equivalent mathematically to a model of the unitary noise described in [33–35]. Quantitatively, one can then describe the evolution of the system by the density operator

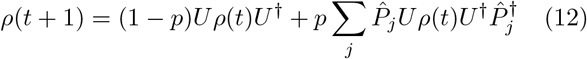

Where 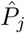 is a projection that represents the action of the decoherence events and *p* is the probability of applying the decoherence per time step. The operator

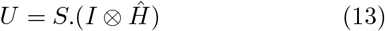

is the unitary operator formed from conditional shift operator

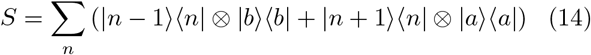

and the Hadamard operator *ĥ*

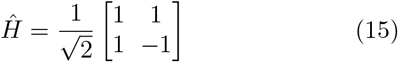

that generates the requisite superposition states of chirality degree of freedom and *I* is the identity matrix [56]. One may therefore also express the evolution of the system as

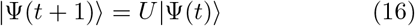

where the state |Ψ(*t*+1) ⟩ is prepared by the superposition of the initial state |*b* ⟩ ⊗ |0 ⟩ by *ĥ* and the conditional translation by the operator *S*. If the Bcd particles in a given chirality state |*j* ⟩ remain localized in the spatial mode *ψ*_*j*_(**r**, *t*), then one can use the approximation

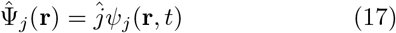

With

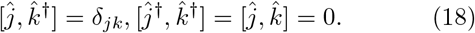

Using these in Eq. (5) above gives

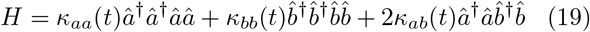

Where

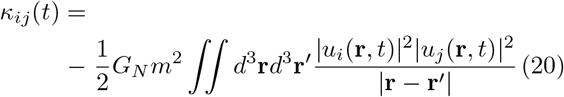

If the wavepackets remain spatially separated by significantly more than their spatial extent for the majority of the evolution, *κ*_*ab*_ *<< κ*_*aa*_, *κ*_*bb*_. Setting *κ*_*aa*_ = *κ*_*bb*_ = *κ*_0_, *κ*_*ab*_ = 0, the unitary operator for the evolution given by Eq. (19) is given by

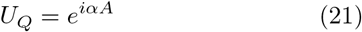

where *α* = *κ*_0_*t/ħ* and 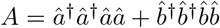. The gravitational interaction between the particles is described by

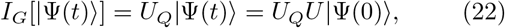

where *U* is now given by Eq. (13). The net effect of all these processes is to generate a non-Gaussian distribution for the Bcd system. Lattice light sheet microscopy studies have revealed highly dynamic Bcd molecules in the posterior end of the embryo with a diffusive-like kinetic behavior [54]. Recent FCS experiments have also measured a significant concentration of Bcd in the posteriormost parts of the embryo [40]. These observations can be explained by analysing the associated probability distribution for the system [33–35]. The probability distribution for the system is given by

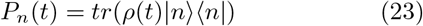

and is calculated by making use of Eq. (12). But one must first define an initial state for the system. Now, the homochirality of amino acids ensures that Bcd, just like all other proteins, is a chiral biomolecule [57]. Since all naturally occurring amino acids that go on to constitute the proteins are left-handed, a reasonable initial state for Bcd will be the one which is dominated by molecules of left-hand chirality configuration: |*b* ⟩⊗ |0 ⟩, where we assume the system to be in *n* = 0 state for particles moving on an integer line. Figure (2) shows the simulated probability distribution of the Bcd gradient for 500 steps of the walker and for numerical values of noise *p* comparable in magnitude to the observed Bcd degradation rates (*k*∼ 10^*−*4^*s*^*−*1^) in the system [58]. Clearly it is steep and non-Guaussian with significant amplitude at both extremes and a vanishingly small amplitude towards the middle. The distribution predicts a small but significant fraction of Bcd on the posterior side of the embryo (right-hand side of Fig. (2)). The underlying physical mechanism, as discussed further below, is then the quantum gravitational interaction between the Bcd molecules.

As with any proposal, including the present one, it is vital, however, that one can distinguish the sought quantum gravitational effect from the analogous effect generated through the electromagnetic interactions. For tests based on optomechanical setups [17], this effect is entanglement, which electromagnetic as well as gravitational interactions would naturally generate. For the test proposed in ref. [15], this is non-Gaussianity which electromagnetic interactions would also generate naturally since they are fundamentally quantum interactions. For the Bcd gradient the concentration, at least in the initial cycles of its formation, is so diffuse (∼ 50 nM [59]) that the provision of any Casimir-Polder or van der Waals forces [60– which vary as ∼ *r*^*−*6^, where *r* is the distance between particles, and which are the only electromagnetic interactions possible here since the Bcd molecules have zero overall charge, is negligible. This can be shown by carrying out a calculation of the mean free path *σ* between the Bcd molecules assumed here as a gas of hard spherical particles. For a gas of hard spheres of mass *m* the expression for mean free path *σ* is given by

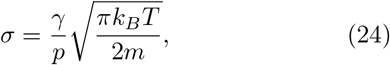

where *γ* is the dynamic viscosity of gas, *k*_*B*_ denotes the Boltzmann constant, *p* the pressure and *T* denotes the absolute temperature [63].

Since the concentration of Bcd protein in the early Drosophila embryo is very small (∼ 50nM [59]), one may take the viscosity of Bcd as similar to the dynamic viscosity of air (∼ 1.849 × 10^*−*5^*kgm*^*−*1^*s*^*−*1^) at room temperature [64] and the pressure *p* as equal to the atmospheric pressure (101.325 × 10^3^ kgm^*−*1^s^*−*2^). The molecular mass *m* of Bcd is ∼9.13 × 10^*−*23^*kg* [37]. Substituting these values in Eq. (24) above yields for the mean free path *σ* at room temperature, *T* = 298.15*K*, a value of ∼ 1.53 ×10^*−*9^m or ~ 1.53nm. At this distance Casimir-Polder or van der Waals forces are expected to be negligible [60–62]. This is because when the intermolecular distances are greater than about 1.0nm the van der Waals force, which varies as ∼*r*^*−*6^, is not strong enough to have any observable consequences. However, the gravitational force, which varies as ∼ *r*^*−*2^, is not negligible at these distances. Furthermore, a key observation from the Bose gas proposal is that, within the Gaussian wave-packet approximation, the minimum number *N* of particles required to observe signatures of quantum gravity is inversely proportional to the particle mass, *N* ∝ 1*/m*, showing that for more massive systems the required number of particles is smaller. For the observed low concentrations of Bcd (about ∼3.4 ×10^8^ Bcd molecules [59]) the van der Waals forces are expected to get reduced even further under such conditions. These factors when taken together are conducive to setting the overall strength of electro-magnetic interactions to zero without much affecting the strength of the sought gravitational interactions. Then a non-Gaussian distribution (see Fig. 2) of Bcd molecules can be attributed only to the gravitational interactions between them, which must therefore be quantum mechanical in origin in the present case.

**FIG 2.**
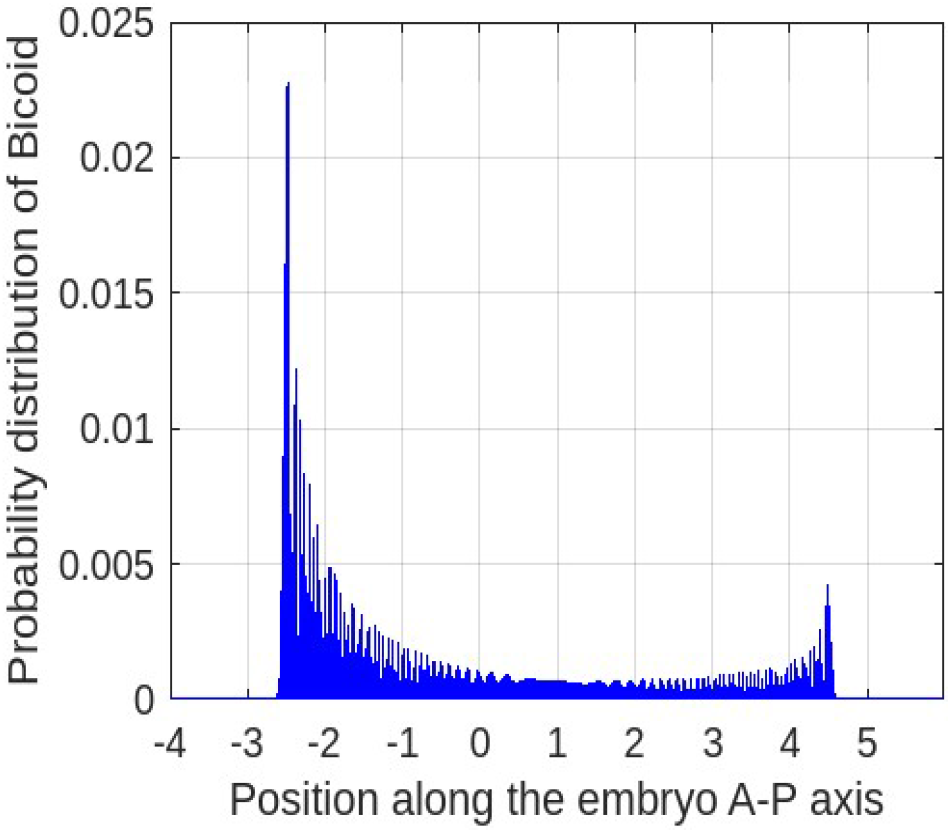
Probability distribution for the quantum-classical model of the Bcd morphogen gradient for 500 steps of the quantum walk predicts a significant concentration of the morphogen at both extremes with a non-Gaussian distribution.

**FIG 3.**
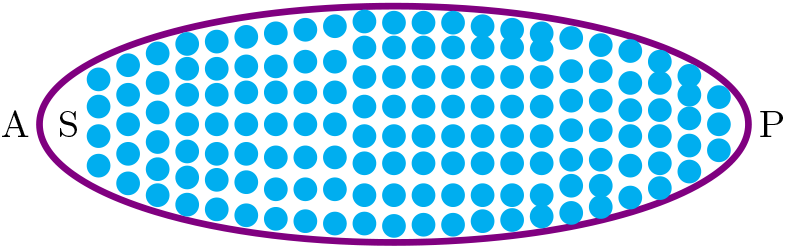
Schematic of the Drosophila embryonic syncytium. The concentration of Bcd inside the fruit fly embryonic syncytium is so dilute (∼ 50 nM) that its dynamic viscosity *γ* and pressure may be approximated by the dynamic viscosity and pressure of air in the atmosphere. A and P denote the anterior and posterior sides of the embryo. Bcd is released from the source located at *S* on the anterior side (A) and diffuses freely towards the posterior side (P) of the syncytium. Thus there is a gradient of Bcd concentration from the anterior to the posterior side. Cyan dots are the nuclei and Bcd can diffuse freely inside the embryonic syncytium.

In this paper we have shown that the phenomenon of Bcd gradient formation in the early fruit fly embryo naturally combines the essential features of two recently proposed experimental tests of quantum gravity, the Bose gas [25] and the BEC proposal [15], in a single quantum system yielding a viable signature of quantum gravity. However, while it has been made clear in the present analysis that the phenomenon of Bcd gradient formation in the early Drosophila embryo is accompanied by quantum gravitational effects, it remains for future experiments to further verify such a noval claim.

I would like to thank Carl Trindle for financial support. Mir Faizal is acknowledged for very informative discussions about quantum gravity and quantum field theory.

## Supplementary Information: A Biological Signature of Quantum Gravity?

### I. TRANSIENT QUANTUM COHERENCES

It is now well-known that fleeting or transient quantum coherences can make some biophysical processes [1–3], like the ones important for quantum biology [4], more efficient by changing the scaling behavior of certain macroscopic observables [5]. However, a more fundamental reason for the prevalence of such short-lived coherences in these processes may be given here. Hamiltonians of actual physical systems, in contrast to their idealized counterparts, typically possess energy mismatches between sites. For such systems long-lived quantum coherences accompanying a diffusion process can lead to a localization problem [6]. Hence only transitory coherences would be preferred in such cases since long-lived coherences in disordered environments can hinder the diffusion of the associated quantum state. Furthermore, there is some evidence for the amplification of coherent quantum states through the constructive action of noise in an otherwise stochastic environment [7].

### II. NON-GAUSSIANITY AS A SIGNATURE OF QUANTUM GRAVITY

For a free and real scalar quantum field 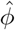 the Hamiltonian contains only the kinetic and mass terms [8] that are necessarily quadratic in the field operators and is given by

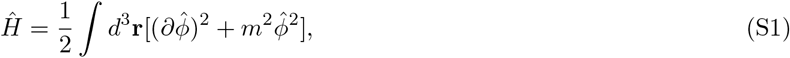

where *m* is the mass of the field and we use the relativistic notation,

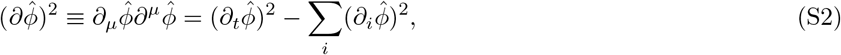

and a Minkowski metric *η*^*µν*^ with signature (+,–,–,–) so that *η*^00^ = +1 [9]. Furthermore,

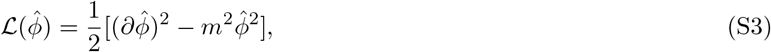

is the corresponding Lagrangian density. Expanding the quantum field in creation and annihilation operators,

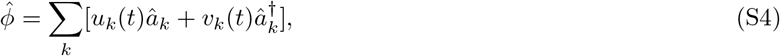

and substituting in Eq. (S1) above results, after collecting terms, in the following Hamiltonian,

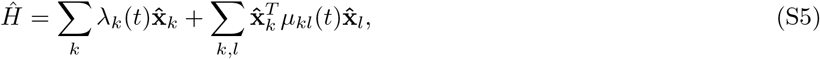

where 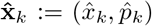 are the quadrature matrices, and *λ*_*k*_(*t*) and *µ*_*kl*_(*t*) are 2 ×1 and 2 ×2 real valued matrices of arbitrary functions of time [8]. Equation (S5) assumes a discrete and finite mode spectrum for simplicity with the extension to infinite and continuous modes being straightforward [10]. Such a Hamiltonian preserves Gaussianity since it is quadratic in field operators [11]. Any other Hamiltonian that is not linear or quadratic in quantum operators will, in general, create non-Gaussianity in the quantum filed state of matter.

If the gravitational field is described by a classical entity 𝒢, dependent on space and time but unable to induce quantum self-interactions of 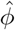, then the interaction of 𝒢with 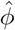 will involve only Hamiltonian terms linear or quadratic in 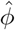 [11]. In other words, the Hamiltonian of this classical interaction would preserve Gaussianity. In contrast, if the gravitational field is represented by a quantum entity 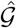, then it is possible for the Hamiltonian to be of order higher than quadratic in the field operators and thus induce non-Gaussianity. Thus, any sign of the creation of non-Gaussianity in the quantum field state of matter would be evidence of quantum gravity. It must be mentioned here that in the non-relativistic Newtonian limit considered here only the temporal component of the perturbed metric is quantized and associated with the Newtonian gravitational potential Φ [12]. In such a limit massive objects interact via the usual 1*/r* Newtonian potential, treated as a quantum operator on the Hilbert space of the masses. Superpositions of the states of massive objects cause superpositions of the metric, which in the non-relativistic limit considered here means superpositions of the Newtonian potential Φ. Such a theory, usually called an effective field theory, is the universal low-energy limit of any theory containing massless spin-2 excitations, called gravitons, and an *S*-matrix [12].

**FIG S1.**
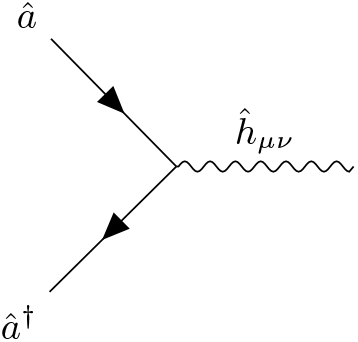
Basic Feynman diagram for the interaction involving quantum gravity where matter emits a graviton, which is associated with *ĥ*_*µν*_. For simplicity, matter is represented by a real scalar quantum field such that *â*^*†*^ and *â* are the creation and annihilation operators of this field. The interaction is then associated with three quantum operators and, therefore, can induce non-Gaussianity.

**FIG S2.**
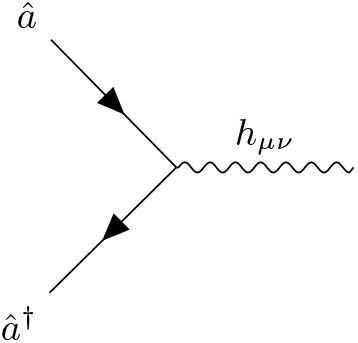
One can illustrate the analogous interaction between matter and classical gravity with a similar diagram except that now the gravitational leg represents a classical gravitational wave *h*_*µν*_ rather than a graviton. Since this classical gravitational interaction is associated with just two quantum operators *â*^*†*^ and *â*, it cannot, in contrast to quantum gravity, induce non-Gaussianity in the quantum field state of matter.

Now, the quantum information can be encoded in discrete variables, such as qubits, or continuous variables, like the eigenmodes of a harmonic oscillator. The continuous variable equivalent of encoding quantum information is applicable to quantum field theory where the fields are considered as collections of harmonic oscillators, thereby allowing one to analyse their information theoretic properties. Application of continuous variable quantum information to quantum gravity shows that a true signature of quantum gravity would be the creation of non-Gaussianity, a continuous-variable resource necessary for universal quantum computation [11]. Non-Gaussianity is generated through non-quadratic quantum operators in the interaction Hamiltonian of a quantum system which in the context of gravity implies that only quantum gravity, compared to classical gravity, can generate it provided other non-gravitational quantum interactions are somehow suppressed. The simplest illustration of this effect in the weak field perturbative regime of gravity is to consider the gravitational interaction between matter to be mediated by gravitons. As illustrated in the figures above, the corresponding Feynman diagrams contain variable numbers of quantum operators and, therefore, may or may not induce non-Gaussianity in the quantum field state of matter [11].

Einstein’s general theory of relativity can be formulated as an action theory with the action *S* decomposed into the Einstein-Hilbert action *S*_*EH*_, containing only gravitational degrees of freedom, and the matter action *S*_*M*_, that tells us how matter and gravity interact:

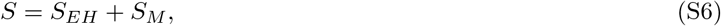

Where

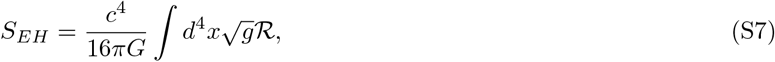

with ℛ the Ricci scalar, and for a real scalar matter field *ϕ*,

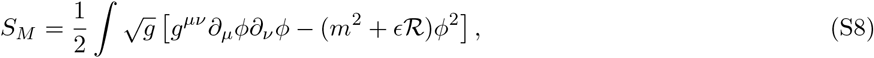

with *ϵ* a numerical factor. The argument presented above for the non-Gaussianity being an indicator of quantum gravity uses only the matter action *S*_*M*_ and says nothing of the purely gravitational action *S*_*EH*_. It relies on the fact that *S*_*M*_ must be, in the absence of all non-gravitational quantum interactions, quadratic in the matter fields such that a classical theory of gravity will preserve Gaussianity. If *S*_*M*_ had terms that coupled gravitational degrees of freedom with nonlinear or non-quadratic functions of the matter fields, then in flat spacetime such terms would still exist and this would result in a new non-gravitational interaction, which we have excluded in our case.

### III. CHAOTIC QUANTUM SYSTEMS & FINE-GRAINED ERGODIC THEOREM

For certain closed or isolated quantum systems the unitary time evolution does not bring the system to a thermodynamic limit after an arbitrarily long time and the systems have to be made ergodic by the addition of a small random perturbation [13]. Then an arbitrary initial state of the system

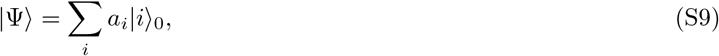

evolving under the action of such a perturbation can be used to obtain a simple formula for the time average of the expectation value of an observable quantity 𝒪 as

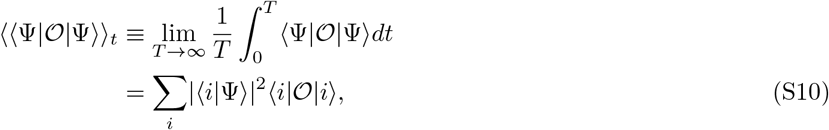

which is known as the fine-grained ergodic theorem [14]. Here *a*_*i*_ is an arbitrary complex amplitude and *T* denotes the absolute temperature. This is unlike the case in a classical analysis, where the notion of ergodicity is easily seen to lead to statistical mechanics.

## IV. LOCC INEQUALITY, SWAPPING & SEPARABILITY

The Bcd system can be considered as a multipartite quantum system with the total Hilbert space ℋ_*B*_ = ℋ_*c*_ ⊗ ℋ_*n*_, where ℋ_*c*_ and ℋ_*n*_ are, respectively, the coin and position Hilbert spaces of the system. Such a system is identical to an ensemble of quantum systems initialised in coherent states following a normal distribution and interacting via a Newtonian gravitational potential [15]. In this multipartite quantum system the global unitary *U* implements a quantum Newtonian Hamiltonian over a gravitationally interacting system initially prepared in random coherent states. For such systems a general and efficiently computable upper bound on the maximal LOCC simulation fidelity has been established recently [15]. These constraints have been dubbed as LOCC inequalities, because they play a role somewhat similar to those of Bell inequalities in the theory of quantum non-locality [16]. Just like the violation of a Bell inequality certifies that the underlying process is unequivocally non-local, in the same way the violation of LOCC inequality certifies that the unitary evolution undergone by the system cannot be a LOCC. In other words, it cannot be explained by introducing a mediator which is a local classical field. These LOCC inequalities can be applied to rule out the LOCCness of a time evolution associated with a gravitational Hamiltonian, even when no entanglement whatsoever is generated at any point in time during the process. Interestingly, as is well-known non-Gaussian states are necessary for the violation of Bell inequalities. An added virtue of above mentioned conceptual step forward is that, while analysing gravitationally interacting quantum systems, it considers an ensemble of possible input states and their resulting output. In this way it better characterises the channel that describes the full dynamics under gravity after a set time. This is a substantial conceptual improvement, because quantum channels, modeling the result of the time evolution to which open quantum systems are subjected, are more general and thus harder to simulate than states. Furthermore, it does not rely at any stage on the creation of gravitationally mediated entanglement.

If the gravitational interaction were to be mediated by a classical field, the time evolution operator it will induce on the system would necessarily be representable by a sequence of local operations assisted by a classical communication channel (LOCC). In such a scheme one is implicitly assuming a very specific meaning for the term *classical gravity* as any gravitational interaction between quantum objects that can be effectively described by LOCC operations. The term *quantum gravity* then refers to any gravitational interaction that cannot be described by LOCC operations. On the other hand, the swap operator comes up naturally when considering a system of gravitationally interacting quantum systems like the one above. For the sake of simplicity, consider the swap operator,

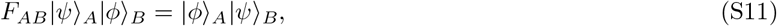

acting on a bipartite system *AB*. This operator is highly non-LOCC as it involves the perfect exchange of quantum information between two parties, which cannot be accomplished with a classical communication channel [17]. In other words, there is no way for the two separate parties Alice and Bob who do not have any access to a quantum communication channel, e.g., an optical fiber, to successfully exchange two unknown quantum states |*ψ* ⟩_*A*_ and |*ϕ* ⟩_*B*_.

Now, an *n*-mode continuous-variable system has Hilbert space ℋ_*n*_ := *L*^2^(ℝ^*n*^). After arranging the dimensionless canonical operators 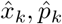 in a vector

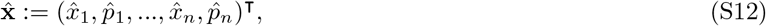

called the modewise decomposition, the canonical commutation relations take the form

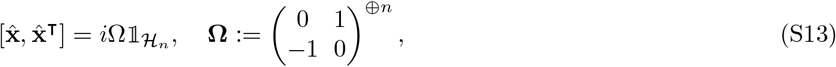

where **Ω** is referred to as the standard symplectic form. It is useful to construct also the annihilation *â*_*j*_ and creation 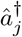 operators, defined by

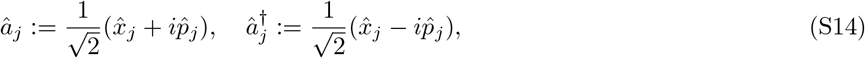

whose commutation relations are written as

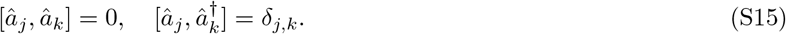

Furthermore, since in the Bcd system the particle number is not fixed but variable, it will be appropriate to invoke the special construction of the Hilbert space called the Fock space,

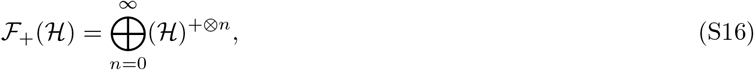

where (ℋ)^+*⊗n*^ is the tensor product of ℋ with itself *n* times and we symmetrize (+) for bosons. Acting repeatedly with creation operators on the vacuum state |0 ⟩, defined via *â*_*j*_|0 ⟩ = 0 for all *j*, yields the Fock states [15]. Other notable states, relevant to us, are the coherent states, parametrized by a complex number *α* ∈ ℂ and given by

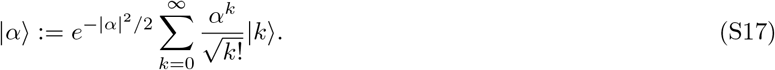

These states form an overcomplete set, in the sense that

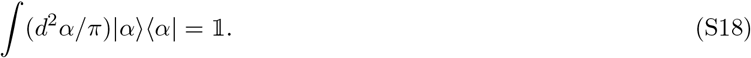

Multimode coherent states are simply product states of single-mode coherent states, i.e.,

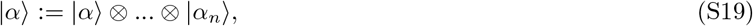

where *α* = (*α*_1_, …, *α*_*n*_)^⊺^ ∈ ℂ^*n*^.

To understand how the above arguments work, we start by looking at our model in which we initially have a system of particles of mass *m* with a free Hamiltonian *H*_0_ given by

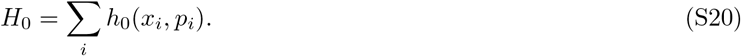

To this picture we would like to add the gravitational interaction, assumed here to be quantum valued, given by

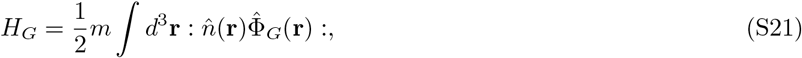

Where

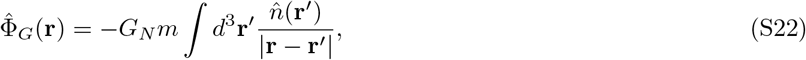

is the quantum valued gravitational potential due to source masses at location **r**^*′*^ [18]. The Hamiltonian that one must basically treat, therefore, is the following

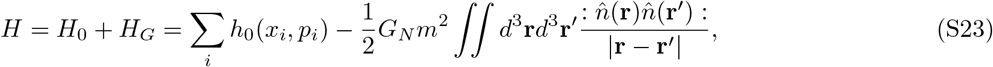

where the gravitational constant is given by *G* = 6.6743 × 10^*−*11^*m*^3^*kg*^*−*1^*s*^*−*2^, as a quantum Hamiltonian acting on the multipartite quantum system. Indeed, in any realistic scenario uncontrolled sources of noise, like the unitary noise in the Bcd system, will always alter the ideal time evolution affected by the Hamiltonian in above equation. Therefore, what one can observe, at best, is some approximate version of Eq. (S23), and intuitively, if the approximation is a sufficiently good one, the impossibility of implementing the dynamics via LOCC should nevertheless hold even in such a case [15]. And indeed, this intuition can be rigorously validated by employing the LOCC inequalities [15]. It must be mentioned here that the Bcd system quantized in ref. [1] is treated as finite dimensional, i.e., 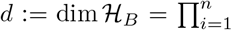. However, as mentioned in the text, in the case of the Bcd system the switch to the continuous and infinite dimensional case is straightforwardly motivated.

Now our multipartite system admits the tensor product structure 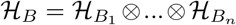. Therefore, we can identify a special class of states on *B* called the fully separable states. A state *ρ*_*B*_ is separable if it admits the decomposition

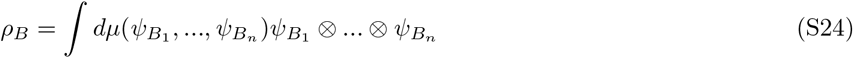

for some Borel probability measure *µ* on the product of the sets of local normalized pure states [15].

The logic behind the LOCC simulation fidelity proceeds as follows [15]. The quantum information-theoretic task that underpins the proposal for detecting the quantum nature of an interaction is that of simulation of a unitary by means of LOCCs. Let *B* = *B*_1_…*B*_*n*_ be our original multipartite quantum system. We would like to ask whether its unitary evolution can be simulated by means by a LOCC as mentioned in the beginning. For this let us consider another, hypothetical, multipartite quantum system denoted 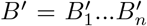. To define the simulation task, we then need two basic ingredients. A source that outputs random pure states of *B* drawn from the ensemble

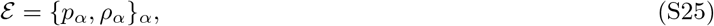

where *ρ*_*α*_ =| *ψ*_*α*_ ⟩⟨*ψ*_*α*_ |, and *p*_*α*_ is the probability for it to be drawn. The index *α* could be discrete or continuous, in which case *p* becomes a probability measure over some measure space [15]. The second ingredient that we need is an isometry

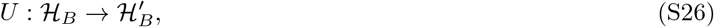

that connects *B* with *B*^*′*^. Both ℰ and *U* are publicly known. At the beginning of each round, a pure state is drawn from ℰ and then handed over to an agent *G*, who knows the ensemble but not its particular realization. *G* then takes one of the following two actions on the system [19].

(1) Null hypothesis (NH): *G* applies *U*.
(2) Alternative hypothesis (AH): *G* attempts to simulate *U* by implementing a LOCC channel Λ ∈ LOCC(*B* → *B*^*′*^).

After *G* has carried out one of these two actions, the output system *B*^*′*^ is sent to a verifier *V*, who knows *α* but not the strategy adopted by *G*. The goal of *V* is to decide between the null hypothesis, corresponding to the pure quantum state 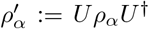,and the alternative hypothesis, corresponding to a possibly mixed state Λ(*ρ*_*α*_). The verifier *V* knows *ρ*_*α*_ and *U*, and hence 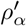, but not the channel Λ. In deciding between (NH) and (AH), *V* can thus make one of two distinct errors.

(1) the type-1 error consists in guessing (AH) while (NH) holds;
(2) conversely, the type-2 error consists in guessing (NH) while (AH) is true.

Knowing both *ρ*_*α*_ and *U*, and thus being able to calculate 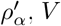 now carries out a quantum measurement represented by the POVM 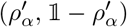 on the unknown state. If the outcome corresponding to 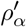 is obtained, *V* guesses (NH), otherwise (AH). In this way, assuming that the measurement is ideal, the probability of a type-1 error is given by

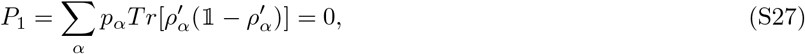

while the corresponding type-2 error probability evaluates to

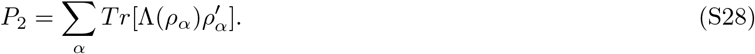

Knowing that *V* will carry out the above test, *G* now attempts to maximize *P*_2_. The maximal value of *P*_2_ that is obtainable by *G* is called the LOCC simulation fidelity (or the classical simulation fidelity) of the isometry *U* on the ensemble ℰ [15]. It is given by

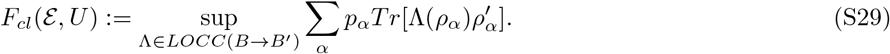

This is nothing but the maximal average fidelity between the target state 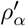 and its simulation Λ(*ρ*_*α*_). In our case *B*_1_, …, *B*_*n*_ are massive quantum systems, Bcd molecules (*m* ∼9.13 ×10^*−*23^*kg*), since they are more than twice the mass of the molecules (*m*∼ 4 ×10^*−*23^*kg*) for which spatial superposition, in the form of matter wave interference, has been observed [20]. *U* is a unitary involving some local terms and an interaction of purely gravitational nature to which above scheme is perfectly applicable. The LOCC simulation of *U* then corresponds to the dynamics induced by a classical (local) gravitational field.

Considering our massive quantum systems *B*_1_, …, *B*_*n*_ subjected to local Hamiltonians *h*_0_(*x*_1_, *p*_1_), …, *h*_0_(*x*_*n*_, *p*_*n*_) and interacting exclusively via gravity (see the main text), as modeled by the Hamiltonian *H*_*G*_, the total Hamiltonian can thus be written as

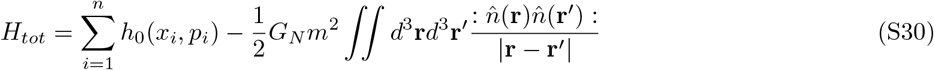

Accordingly, the time-evolution operator associated to some time interval *t* is represented by the unitary

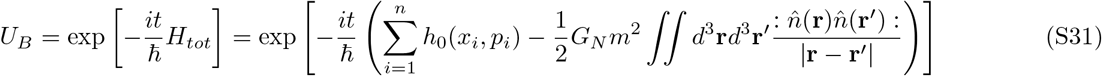

acting on *B*. Suppose now that for our multipartite system *B*, unitary *U*_*B*_ as in Eq. (S31), and ensemble 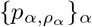,we have computed the LOCC simulation fidelity *F*_*cl*_(ℰ, *U*) in Eq. (S29). One now imagines to run the simulation experiment discussed above many times independently, drawing the initial pure states at random in an i.i.d. fashion. If gravity behaves as a quantum Hamiltonian, then in the case of ideal measurements we will always get the outcome (NH) corresponding to a null hypothesis guess. On the contrary, if gravity behaves as a local classical field, then one will get the outcome (AH) corresponding to an LOCC simulation guess with frequency at least 1-*F*_*cl*_(ℰ, *U*). That is to say if one obtains (AH) with frequency lower than 1-*F*_*cl*_(ℰ,, *U*), one can conclude that gravity did not behave as a local classical field. It turns out that an exact computation of 1-*F*_*cl*_(ℰ,, *U*) is not needed; even an upper bound such as

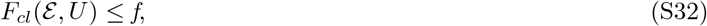

where *f*∈[0, 1), suffices because it enables one to state that obtaining (AH) with frequency lower than 1-*f*, [and hence of 1− *F*_*cl*_(ℰ,, *U*)] is direct evidence of the quantum nature (i.e., non-LOCCness) of the gravitational interaction. In this sense, the status of the inequality in Eq. (S32) is analogous to that of a Bell inequality in the theory of nonlocality [16]. Just like an experimental violation of Bell inequality provides a definitive proof that the underlying process producing the correlations is nonlocal, in a similar way the experimental violation of above inequality would prove that the dynamics undergone by the system does not conform to any LOCC model. In light of this reasoning, the inequality above is referred to as the LOCC inequality [15].

## Notes

### Competing Interest Statement

The authors have declared no competing interest.

